# DNA methylation and proteomics integration uncover dose-dependent group and individual responses to exercise in human skeletal muscle

**DOI:** 10.1101/2022.07.11.499662

**Authors:** Macsue Jacques, Shanie Landen, Javier Alvarez Romero, Danielle Hiam, Ralf B. Schittenhelm, Iresha Hanchapola, Anup D. Shah, Nir Eynon

**Affiliations:** Institute for Health and Sport (iHeS), Victoria University, Melbourne, Australia; Institute of nutrition and health sciences, Deakin University, Melbourne, Australia; Monash Proteomics & Metabolomics Facility, Monash University, Melbourne, Australia

**Author notes:** Corresponding Author: Professor Nir Eynon, Institute for Health and Sport (iHeS), Victoria University, PO Box 14428, Melbourne, VIC 8001, Australia.

**Keywords:** Exercise metabolism, skeletal muscle, epigenetics, DNA methylation, proteomic

## Abstract

**Objective:** Exercise is a major regulator of muscle metabolism, and health benefits acquired by exercise are a result of molecular shifts occurring across multiple OMIC levels (i.e. epigenome, transcriptome, proteome). Identifying robust targets associated with exercise response, at both group and individual levels, is therefore important to develop health guidelines and targeted health interventions.

**Methods:** Twenty, apparently healthy, moderately trained (VO_2_ max= 51.0±10.6 mL·min^−1^·kg^−1^) males (age range= 18-45yrs) from the Gene SMART (Skeletal Muscle Adaptive Responses to Training) study completed a 12-week High-Intensity Interval Training (HIIT) intervention. Muscle biopsies were collected at baseline and after 4, 8, and 12 weeks of HIIT. High throughput DNA methylation (∼850 CpG sites), and proteomic (∼3000 proteins) analyses were conducted at all-time points. Mixed-models were applied to estimate group and individual changes, and methylome and proteome integration was conducted using a holistic multilevel approach with the mixOmics package.

**Results:** Significant shifts in the methylome (residual analysis) and proteome profiles were observed after 12 weeks of HIIT. 461 proteins significantly changed over time (at 4, 8, and 12 weeks), whilst only one differentially methylated position (DMP) was changed (adj.p-value <0.05). K-means analysis revealed clear protein clustering exhibiting similar changes over time. Individual responses to training were observed in 101 proteins. Seven proteins had a large effect-sizes >0.5, among them are two novel exercise-related proteins, *LYRM7* and *EPN1*. Integration analysis uncovered bidirectional relationships between the methylome and proteome.

**Conclusions:** We showed a significant influence of HIIT on the epigenome and proteome in human muscle, and uncovered groups of proteins clustering according to similar patterns across the exercise intervention. Individual responses to exercise were observed in the proteome with novel mitochondrial and metabolic proteins consistently changed across individuals. Future work is required to elucidate the role of such proteins in response to exercise as well as to investigate the mechanisms associating genes and proteins in response to exercise.

## 1. Introduction

Exercise training is strongly associated with health benefits such as increased oxygen consumption, and significant reduction in the risk for chronic diseases (e.g., type 2 diabetes, cardiovascular disease) [1-4]. These benefits are elicited as a result of exercise-induced changes in genes and proteins, facilitating a positive shift in cell signalling and metabolism [5-9]. Identifying exercise-enhanced molecules is crucial for the development of targeted therapies to treat metabolic diseases and/or provide guidelines for future health policies.

High throughput unbiased OMIC analyses, including genomic, epigenomic, transcriptomics and proteomics, have become more readily available and are increasingly used to robustly identify exercise marks in healthy and diseased populations [10; 11]. While genomic and transcriptomic control of exercise adaptations have been well studied in both athletes and the general population [11-14], epigenetic [15-18] and untargeted proteomic analyses [19; 20] are fairly new in this field of research [17; 21]. Epigenetic mechanisms are highly sensitive to environmental stimuli and can affect how genes are transcribed. Perhaps the most important and studied epigenetic modification is DNA methylation [22]. It involves the addition of a methyl group (CH3) to cytosines in the DNA [23]. DNA methylation levels are altered in several diseases including cancer [23] and metabolic syndrome [24]. Research into exercise training and epigenetics is still in its early stages, with few studies investigating DNA methylation changes after exercise training [17; 18; 25; 26]. These studies suggest that exercise training may induce changes in the methylation status of genes involved in muscle function. Thus, it may be possible to model a long-lasting advantageous gene and protein expression patterns for superior responses to exercise [7; 15; 17; 18], although it should be acknowledge that causality studies are lacking. Furthermore, it is currently unknown whether exercise-induced changes in these methylation loci are associated with changes in proteins.

The majority of the cellular processes are controlled by proteins, and utilising high throughput proteomic analyses offers great potential in investigating molecular mechanisms underlying exercise-induced adaptations [8]. Proteins are the end product of genetic, epigenetic, and (post-) translational processes, and may be regarded as the interface between the genome and the environment [27]. For example, recent time series analyses on multiple omics have revealed several proteins belonging to four different clusters (including proteins involved in inflammatory response, immune response, oxidative stress, etc.) to significantly change post-exercise over time in blood [6]. Long-term exercise adaptations are characterised by a combination of acute exercise bouts, however only a handful of studies have investigated changes in the proteome in response to long term exercise in skeletal muscle [6; 27; 28].

Personalised medicine is a hot topic in exercise studies. However, individuals respond differently to similar exercise programs, and there is no ‘one size fits all’ approach that will benefit everyone [4; 29-31]. To address the issue of inter-individual variability (i.e. variability observed between individuals), the study design should also incorporate intra-individual variation (i.e. variability observed within same individual – individual response to training) [32]. To date, several methods have been proposed to study individual response to training [31-35], however as yet most studies only assess exercise responses at the group level, and only few have applied a suitable protocol to appropriately uncover the individual response to exercise [34; 36-38], with no study exploring individual responses in large omics data.

Here, we have utilized the repeated testing approach, and OMIC integration (which has been explored and explained in detail elsewhere [34; 38; 39]) to uncover robust exercise-induced time-course changes (at baseline, 4 weeks, 8 weeks, and 12 weeks) in the epigenome (DNA methylation) and proteome profiles in human skeletal muscle.

## 2. Methods

### 2.1 Participants

Twenty participants from the Gene SMART (Skeletal Muscle Adaptive Responses to Training) study [40] commenced the study, and 16 participants completed 12-week of **High-Intensity Interval Training** (HIIT). From the 20 initially recruited participants, **19 completed 4 weeks of HIIT** (1 dropout), **of these 18 completed 8 weeks** (1 dropout), **and 16 completed the full 12-weeks of HIIT** (1 dropout and 1 exclusion due to inconsistent results (i.e. duplicate tests provided more than 10% difference).

Participants were apparently healthy, moderately trained males (VO_2max_ 51.0±10.6 mL·min^−1^·kg^−1^), aged 18 to 45 years old. The study was approved by the Victoria University human ethics committee (HRE13-223 & HRE21-122) and written informed consent was obtained from each participant. Participants were excluded from the study if they had a past history of definite or possible coronary heart disease, significant chronic or recurrent respiratory condition, significant neuromuscular, major musculoskeletal problems interfering with ability to cycle, uncontrolled endocrine and metabolic disorders or diabetes requiring insulin and other therapies [40].

### 2.2 Study design

Participants were tested at baseline and after 4 weeks, 8 weeks, and 12 weeks of HIIT. To ensure progression, training intensity was re-adjusted every 4 weeks based on the newly determined Peak Power output (W_peak_) and Lactate Threshold (LT) from the Graded Exercise Test (GXTs). These tests also allowed monitoring individual progress of participants for the longitudinal analysis of training adaptations. To increase accuracy in measurement and to reduce biological day-to-day variability in participants’ performance, physiological measures of fitness (W_peak_, LT, and VO_2max_) were assessed from two GXTs conducted at each time point **(Figure 1**).

**Figure 1:**
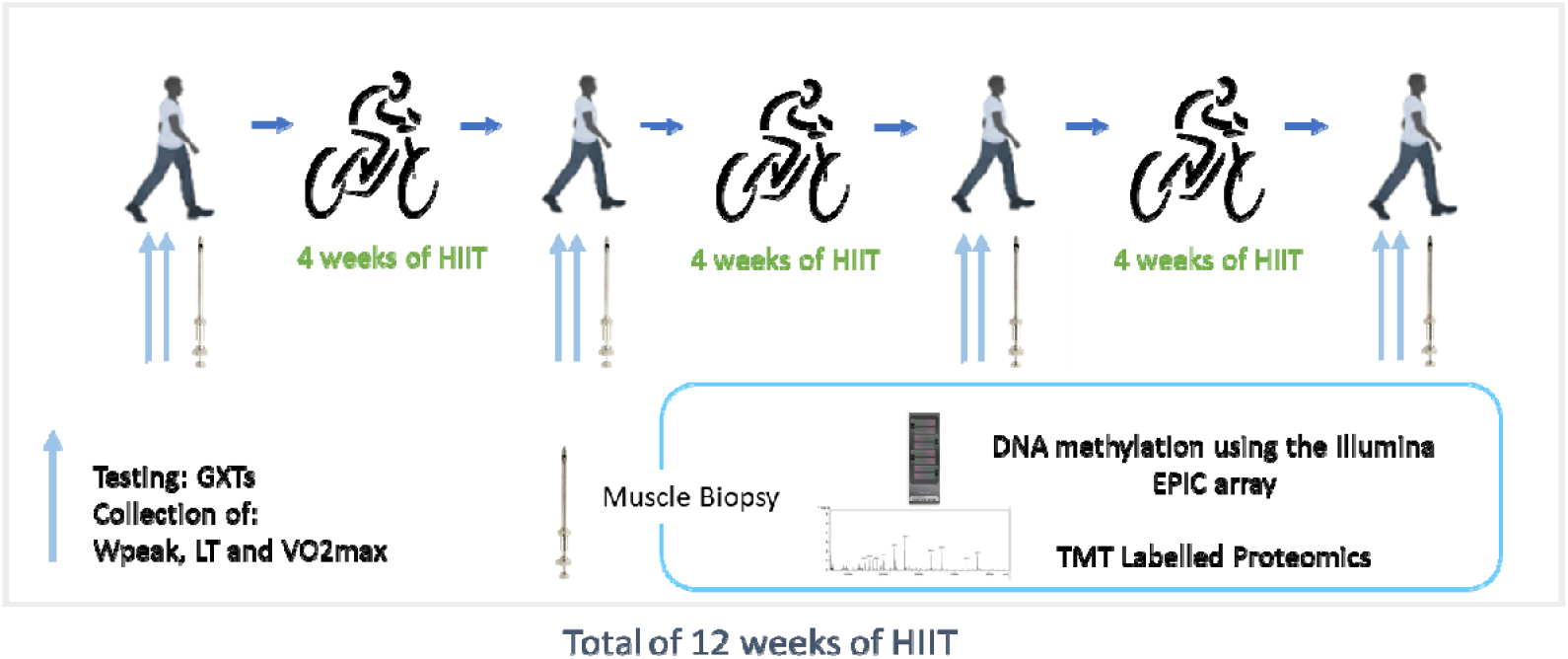
Study design

### 2.3 Muscle biopsies

A controlled diet for 48 h prior to the muscle biopsies was provided to the participants, according to the guidelines of the Australian National Health & Medical Research Council (NHMRC). Muscle biopsies were taken by an experienced medical doctor using a Bergstrom needle from the vastus lateralis muscle of the participants’ dominant leg, following local anaesthesia (2mL, 1% Lidocaine (Lignocaine)). The needle was inserted in the participant leg and manual suction was applied for muscle collection. Excess blood from biopsy was removed using an absorbent sheet, and muscle was then immediately frozen in liquid nitrogen and stored at -80 degrees until further analyses. Muscle biopsies were collected at 4 timepoints (Pre, 4W, 8W and 12W) for comprehensive analyses of DNA methylation and proteome (**Figure 1**).

### 2.4 DNA extraction and DNA methylation analyses

DNA was extracted using the Qiagen All prep DNA/RNA kit (Cat/ID: 80204), according to manufacturer guidelines. In brief, ∼15mg of muscle samples were homogenised and separated by a column into genomic DNA and RNA, and then eluted separately. RNA was stored in -80 degrees for future analyses while dsDNA concentrations were estimated with the Invitrogen Qubit Fluorometer. DNA methylation analyses was carried by the Illumina Infinium Methylation EPIC array (https://www.illumina.com/products/by-type/microarray-kits/infinium-methylation-epic.html) according to manufacturer protocols. Briefly, the DNA input amount was 500ng for bisulfite conversion. QC of bisulfite conversion was carried by MPS (Methylation specific PCR) on specific regions. Once QC was evaluated, whole genome amplification was conducted followed by array hybridization, single base extension before the array was scanned.

### 2.5 Pre-Processing and Statistical Analyses

Pre-processing of the DNA methylation data was conducted, using the ChAMP analysis pipeline [41] in the R statistical software (www.r□project.org). Data was imported into R as IDAT file. Probes were filtered based on detection p-value >0.01 (default value for the champ.load function) and were filtered out if <3 beads in at least 5% of samples, missing β□values, non-CpG probes or if probes aligned to multiple locations in the data set. Additionally, SNP-related probes (“EUR” population probes described in Zhou et al [42]) and probes located within sex chromosomes (this was performed as the full dataset contained males and females, but here we are only using the male subset data) were filtered out. β - values were then obtained as follows:

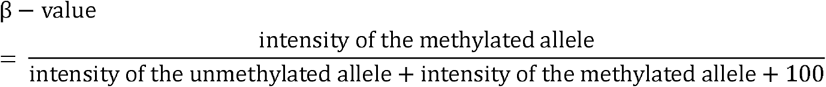

A β□mixture quantile normalisation method was conducted to normalize the data for Type I and Type II probes, generated from the HMEPIC array. We then performed a singular value decomposition (SVD) analyses to identify variations in each individual dataset. Because chip position as well as batch were significantly identified as sources of variation, data was converted to M-values and then normalized for batch and chip position using the ComBat function in the sva package. Finally, a final quality check (QC) was performed in the data to ensure all sources of unwanted variability were accounted for.

Following normalisation and QC procedures, we undertook differential methylation analysis by converting β-values to M-values since those present distributions that are more appropriate for statistical test and differential analysis of methylation levels [43]:

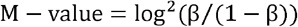

To identify exercise induced timeline changes in DNA methylation we used linear mixed models using the lmerTest [44] package. The models were built as follows:

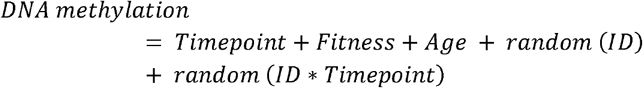

Where outcome was the DNA methylation probes, timepoint (PRE, 4WP, 8WP and 12WP), covariates in this case was age and baseline fitness, and ID was individual identification for each participant. DMPs associated with either fixed (group level) or random (individual level) effects were considered significant if adjusted p-value <0.05. If significant DMPs were identified we proceed the analyses by identifying if such positions were associated with specific regions (DMRs: clusters of DMPs that presented consistent DNA methylation change with fitness measure), for this we used the DMRcate [45] package. To identify fitness-associated pathways, we performed a gene set enrichment analysis using two methods. 1-The champ.ebGSEA function which does not consider pre-determined significance of DMPs, but instead performs its own analyses across multiple genes identified by DNA methylation probes and phenotype selected. 2-The GOmeth methodology which requires pre-specified significant positions [46]. All pathways with an adjusted p-value <0.05 were considered significant and p-values obtained from both analyses were adjusted using the Benjamini & Hochberg method (BH).

Plots were created utilizing the following packages: ggplot2 [47], ggpubr [48], complexHeatmaps [49], FactorMiner [50]. All analyses were performed using R software version 4.0.2.

#### 2.5.1 Proteomics extraction and analyses

Muscle tissue was lysed in 300 ul SDS solubilization buffer (5% SDS, 50 mM TEAB, pH 7.55), heated at 95°C for 10 min and then probe-sonicated before measuring the protein concentration using the BCA method. A total protein amount of 100µg (suspended in 50µl) was used for each sample for subsequent analyses. The lysed samples were denatured and alkylated by adding TCEP (Tris(2-carboxyethyl) phosphine hydrochloride) and CAA (2-Chloroacetamide) to a final concentration of 10mM and 40mM, respectively, and the mixture was incubated at 55°C for 15 min. Sequencing grade trypsin was added at an enzyme to protein ratio of 1:50 and incubated overnight at 37°C after the proteins were trapped using S-Trap mini columns (Profiti). Tryptic peptides were eluted from the columns using (i) 50 mM TEAB, (ii) 0.2% formic acid and (iii) 50% acetonitrile, 0.2% formic acid. The fractions were pooled, concentrated in a vacuum concentrator and reconstituted in 40µl 200 mM HEPES, pH 8.5. Using a Pierce Quantitative Colorimetric Peptide Assay Kit (Thermo Scientific), equal amounts of peptides for each sample were labelled with the TMTpro 16plex reagent set (Thermo Scientific) according to the manufacturer’s instructions, this labelling strategy was implemented to minimise channel leakage. Individual samples were then pooled and high-pH RP-HPLC was used to fractionate each pool into 12 fractions, and were acquired individually by LC-MS/MS to maximise the number of peptide and protein identifications.

Using a Dionex UltiMate 3000 RSLCnano system equipped with a Dionex UltiMate 3000 RS autosampler, an Acclaim PepMap RSLC analytical column (75µm x 50cm, nanoViper, C18, 2µm, 100Å; Thermo Scientific) and an Acclaim PepMap 100 trap column (100µm x 2cm, nanoViper, C18, 5µm, 100Å; Thermo Scientific), the tryptic peptides were separated by increasing concentrations of 80% acetonitrile (ACN) / 0.1% formic acid at a flow of 250nl/min for 158min and analyzed with an Orbitrap Fusion Tribrid mass spectrometer (ThermoFisher Scientific). The instrument was operated in data-dependent acquisition mode to automatically switch between full scan ms1 (in Orbitrap), ms2 (in ion trap) and ms3 (in Orbitrap) acquisition. Each survey full scan (380–1580m/z) was acquired with a resolution of 120,000, an AGC (automatic gain control) target of 50%, and a maximum injection time of 50ms. Dynamic exclusion was set to 60 seconds after one occurrence. Keeping the cycle time fixed at 2.5sec, the most intense multiply charged ions (z ≥ 2) were selected for ms2/ms3 analysis. Ms2 analysis used CID fragmentation (fixed collision energy mode, 30% CID Collision Engery) with a maximum injection time of 150ms, a “rapid” scan rate and an AGC target of 40%. Following the acquisition of each MS2 spectrum, an ms3 spectrum has been acquired from multiple ms2 fragment ions using Synchronous Precursor Selection. The ms3 scan was acquired in the Orbitrap after HCD collision with a resolution of 50,000 and a maximum injection time of 250ms.

The raw data files were analyzed with Proteome Discoverer (Thermo Scientific) to obtain quantitative ms3 reporter ion intensities

#### 2.5.2 Proteomics pre-processing and analysis in the Gene SMART cohort

A total of 3368 proteins were identified in this study. The data pre-processing and downstream statistical analysis was performed using the R statistical software (www.r□project.org). Before normalisation, proteomic data was filtered for high confidence protein observations and contaminants and proteins that were only identified by a single peptide or proteins not identified/quantified consistently across the experiment were removed. Remaining missing values were imputed using MNAR method, assuming the missing values were due to low expression for such proteins, and then normalised using the VSN method. Both imputations and VSN were conducted in the *DEP* package [51]. Batch effects were normalised with the internal referencing scaling (IRS) method [52] by the use of reference channels. After normalisation steps 2365 proteins were included in subsequent analyses.

Pre-processed proteomic data (described above) was used in this analyses. Protein expression (logFC) were regressed against baseline fitness (VO_2max_), Age and timepoint (PRE, 4WP, 8WP and 12WP) as covariates:

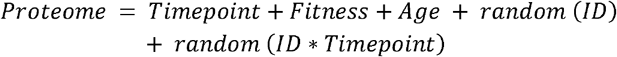

Results with an FDR <0.05 were deemed significant.

### 2.6 PCR validations

Based on our proteomic results we have then selected the top proteins presenting the largest effect sizes and that based on the literature could be of interest to further explore in the exercise context to perform transcriptome analysis to investigate if protein expression aligned with transcript expression at the same timepoints.

RT-PCR for USP2, AMPD3 and SOD3 was performed on muscle. RNA was extracted using the AllPrep DNA/RNA FFPE Kit (#80234 Qiagen). 10 ng of RNA was then diluted into 50ul and reverse transcription was conducted using the iScript™ Reverse Transcription Supermix for RT-qPCR (Bio-Rad) with a thermomixer. Primers for USP2, AMPD3 and SOD3 used for this experiment are described below (**Table 1**). RT-PCR was conducted using the QuantumStudio-7 (Bio-Rad). mRNA expression levels were quantified by real-time PCT using SYBR green fluorescence. Cycle threshold (Ct) values were normalized to a housekeeping gene, Cycl1. Samples were analysed in triplicates and data was manually curated. In cases where samples yielded a standard deviation > 0.4, the divergent sample was removed.

**Table 1:**
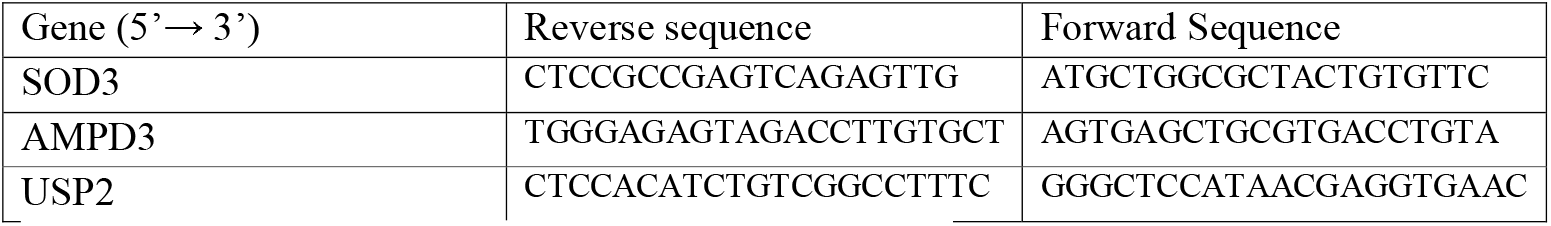
Primer information for each gene used in the validation step.

### 2.7 Integration analysis

Integration between DNA methylation and Proteome data was performed using a holistic approach where the 10,000 closest to significance CpGs and all identified proteins were included in the analysis. Integration was conducted with the mixOmics package, using a multilevel approach to account for repeated measures [53; 54].

## 3. Results

### 3.1 DNA Methylation and proteomic profiles change over the course of a 12-week exercise intervention

Firstly, we investigated if exercise triggered dose-response changes (baseline, 4 weeks, 8 weeks, 12 weeks) in the methylome and proteome (**Figure 1**). Changes in physiological and mitochondrial markers during this intervention are described in detail elsewhere [37; 38]. After adjusting for multiple testing, only one DNA methylation loci (DMP) changed significantly over time (cg23669611, adj.p-value=0.043). However, there was an inflection of p-values towards zero (**Figure 2A**), suggesting that changes in the methylome are occurring but did not reach statistical significance, potentially due to a small sample size. Given this inflation, we have further analysed our data by removing the time variable from the mixed model and extracted the residuals from the 10,000 closest to significance CpGs and performed a PCA analysis (**Figure 2B**). Based on Figure 2B, CpG with similar methylation levels are located closer together on the PCA plot. A significant shift in the methylome towards the left on dimension one (dim.1) occurred as training intervention progressed (p-value=0.0000352).

**Figure 2:**
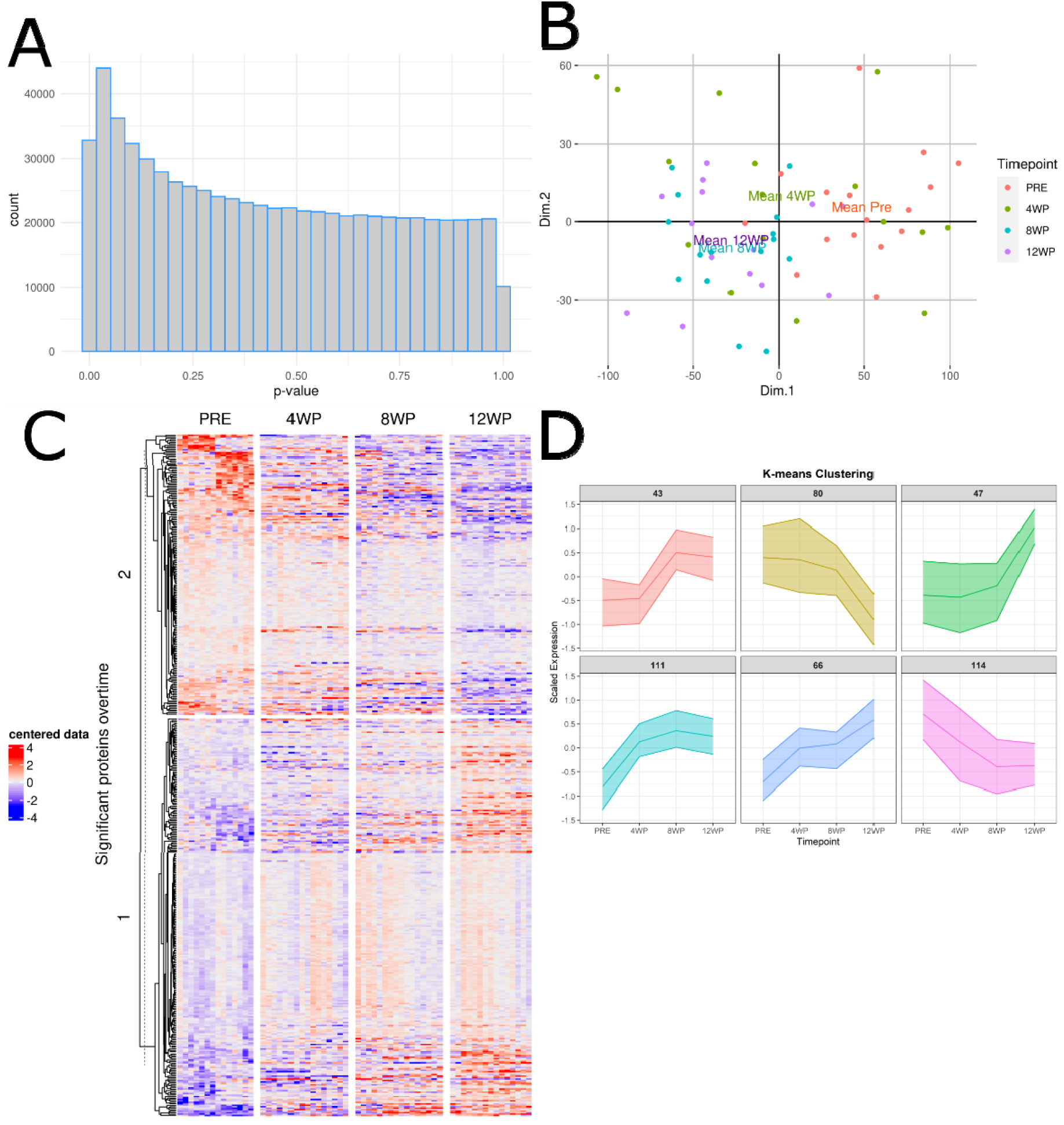
DNA methylation and proteomics changes over the course of the exercise intervention. A: Histogram of inflation of p-values based on DNA methylation results from mixed model, and x-axis represent number of CpGs. B: PCA plot of DNA methylation residuals change over time. Means are highlighted by text. Significant changes over time was observed in the methylome (Estimate: -26.151, SD:5.108, p.value=0.0000352). C: Heatmap of significant proteins changes over time in the group level (n = 461, p < 0.05). D: K-means plot representing trajectory change of significant proteins over time. Each cluster is represented by a group of proteins that present similar changes during the exercise intervention. The number on top of each individual plot represents the number of proteins that belong to each cluster. For example in cluster 1 out of the 461 significant proteins 43 showed positive changes between 4 weeks and 8 weeks.

Of the 2635 proteins included in the analysis 461 proteins significantly changed across the training intervention (**Figure 2C & Supplementary Table 1**). There were 26 proteins that are involved with NADH dehydrogenase, 10 with ATP synthase and 18 were mitochondrial ribosomal proteins. Interestingly, unlike mitochondrial ribosomal proteins, nuclear ribosomal protein expression decreased over time, concomitant with a decrease in the expression of proteins involved in eukaryotic translation. Next, we assessed whether the 461 significantly changing proteins exhibit specific trends over time (i.e. proteins that changed after 4 weeks, or after 8 weeks, etc). We performed k-means analyses with the 461 proteins which were divided into 6 clusters **(Figure 2D & Supplementary Table 2)**. This clustering revealed coordinated timeline changes with some proteins increasing only between 4WP to 8WP (Cluster 1) while others increased only between 8WP to 12WP (Cluster 2). Over-representation analyses (ORA) of all DE proteins identified 14 significant pathways (adj.p-value <0.05). Interestingly, all 14 pathways were related to mitochondrial function (**Figure 3A**). When ORA was performed to each individual cluster of proteins (**Figure 2D**), only cluster 4 and 5 resulted in significant pathways (adj.p-value <0.05). Proteins belonging to cluster 4 exhibited a pattern of increase in protein expression up to 8 weeks of training before plateauing in the last 4 weeks. This cluster resulted in 18 significant pathways of which most were linked to mitochondrial functioning (**Figure 3B**). Similarly cluster 5 presented increases in protein expression throughout the intervention with smaller increases observed between 4 to 8 weeks. Pathways identified for this cluster (n=6) were also linked to mitochondrial function and metabolism (**Figure 3C**). All other individual clusters did not result in any significant pathways.

**Figure 3:**
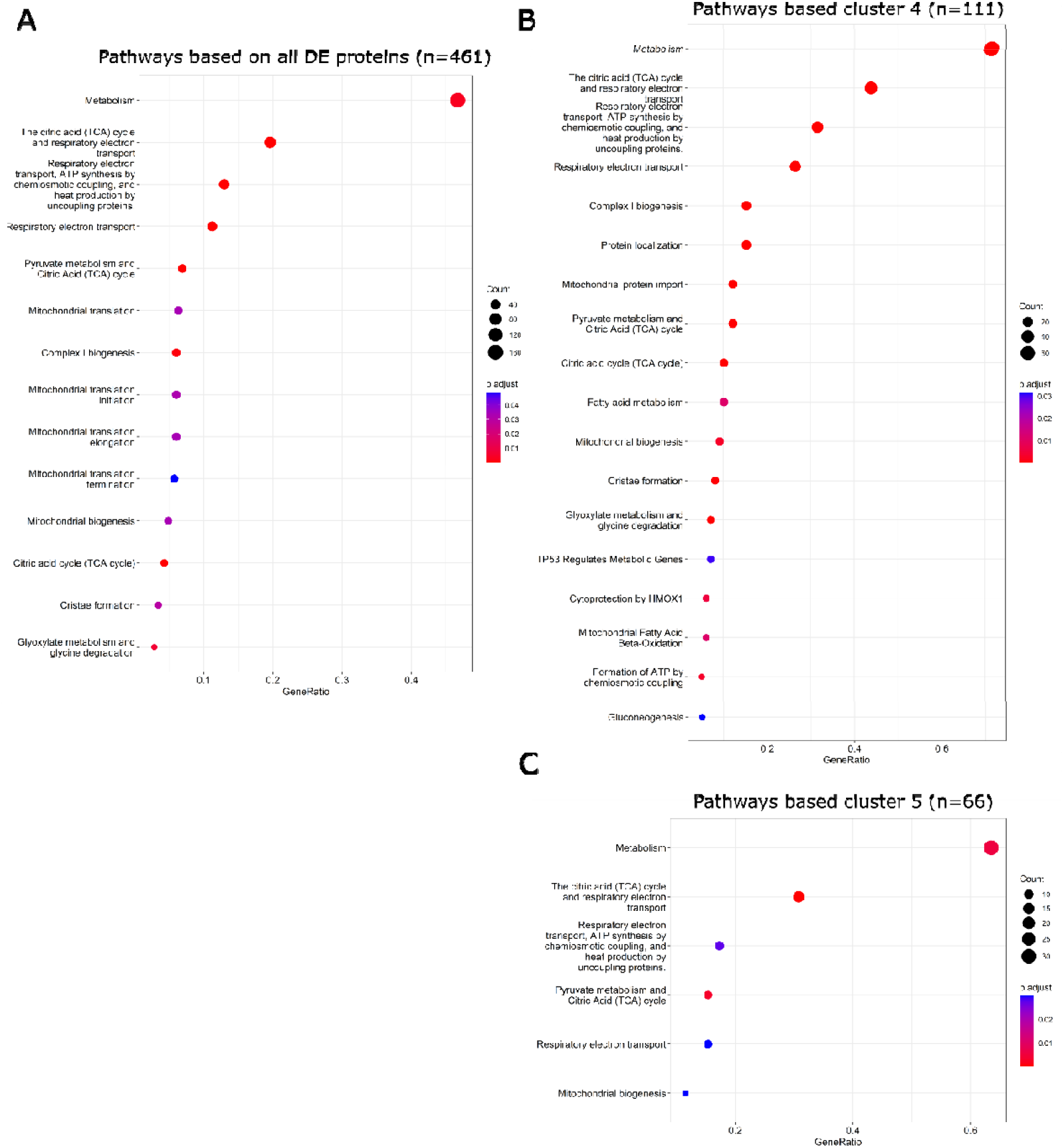
Over-representation enrichment analyses of DE proteins and protein clusters. A: Dotplot of significant reactome pathways for all DE proteins (n=461). B: Dotplot of significant reactome pathways for proteins cluster 4 (n=111). C: Dotplot of significant reactome pathways for proteins cluster 5 (n=66). In A, B and C the size of the dots represents the number of genes assigned to each pathway and the colour of dots represents significance based on adjusted p-value of <0.05.

### 3.2 Individual responses in DNA methylation and proteome over the course of the exercise intervention

We utilised a repeated testing approach as a proxy to estimate variability observed around individual slopes, which allowed us to estimate with more certainty individual changes in response to training. One CpG (cg06587054, adj.p-value=0.001), and 101 proteins have shown consistent individual responses over time (**Figure 4A and Supplementary Table 3**), of which 52 overlapped with group level time course response. The significant CpG was located on the *RPSAP31*;*PARL* protein-coding gene, and *PARL* is also known as a mitochondrial rhomboid protease [55]. Of the significant proteins, 7 had a large effect sizes >0.5 (**Figure 4B**), including: USP2, a protein known to be involved in biological rhythms (i.e. circadian clock), and ageing [56], and differentiation of myoblasts to myotubes [57]. TRIM28, a chromatin regulator that mediates repression of many transcriptional factors [58]. SOD3, an antioxidant enzyme that catalyses the conversion of superoxide radicals, protecting tissues from oxidative stress [58]. RHOT1, a mitochondrial GTPase involved in mitochondrial trafficking [58]. LYRM7 which works as a chaperone of the inner mitochondrial complex III (the main enzyme complex in the mitochondrial respiratory chain) stabilizing this matrix prior to its translocation and insertion into the late CIII dimeric intermediate within the mitochondrial inner membrane [58]. EPN1 which is involved in endocytosis, and loss of function of this gene is associated with reduced tumour growth and progression of cancer [58], and, AMPD3 which has a critical role in balancing energy charge and nucleotide metabolism [59; 60]. While most of these proteins have been previously reported to be associated with either exercise [26; 61-67] and/or skeletal muscle [26; 63-65; 68], SOD3 and RHOT1 have not been previously studied in skeletal muscle. TRIM28 is novel in the exercise context, whilst EPN1 and LYRM7 have not been previously associated with either skeletal muscle or exercise response. Finally, when considering all 101 proteins significant for trainability no significant pathways (adj.p-value <0.05) were identified after ORA analyses.

**Figure 4:**
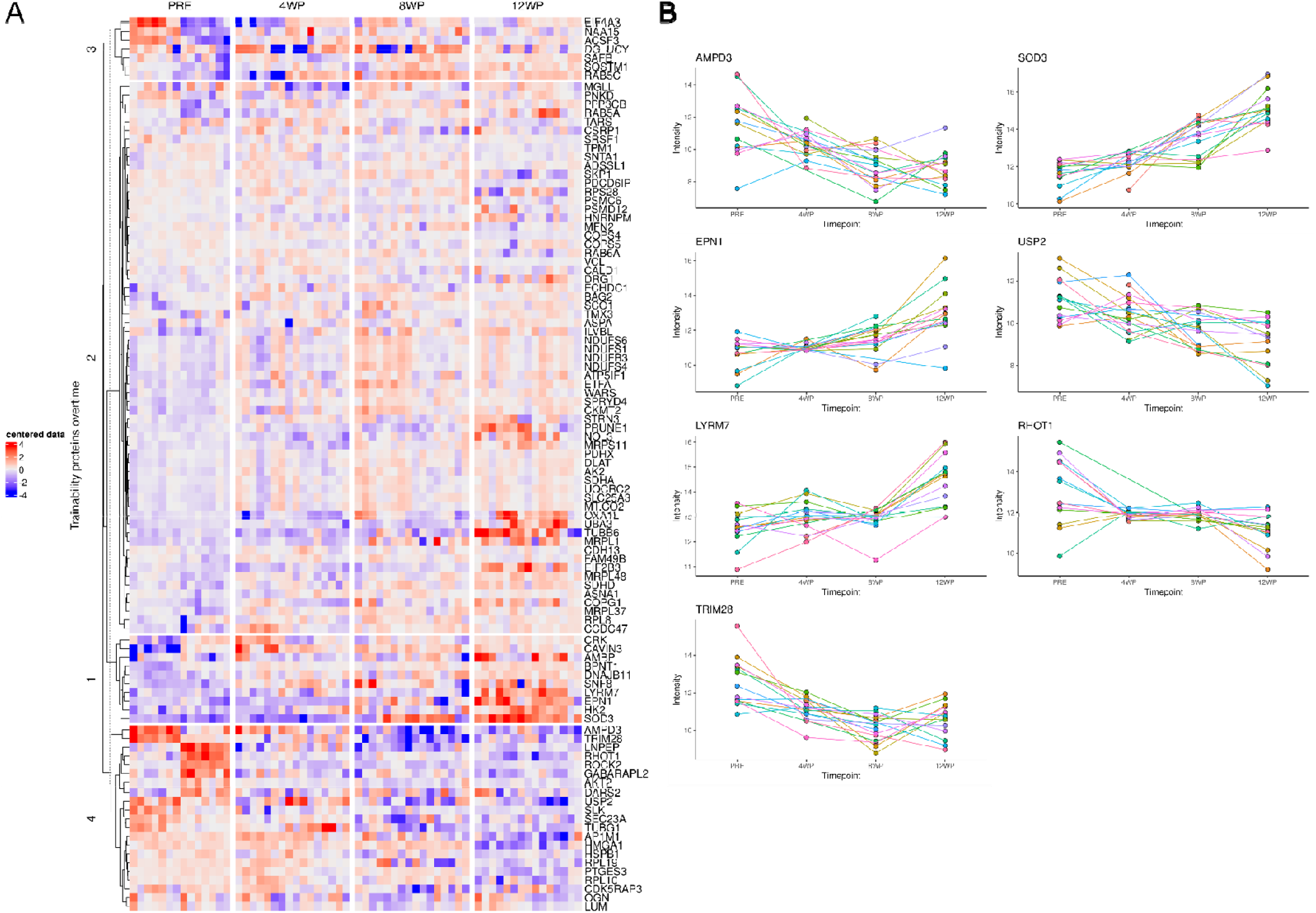
Trainability Proteins. A: Heatmap with the 101 proteins where individual training response was estimated as significant (p<0.05). B: Out of the 101 proteins 7 have shown a cha in effect size >0.5 and are represented by a scatterplot with individual points for each participant. Each participant is represented in a different colour and colour is unique to participant.

### 3.3 Integration of DNA methylation and proteome based on individual response and transcriptome validation

Given the lack of power to detect DNA methylation changes when all the CpGs derived from the EPIC array are included, we performed a filtering step based on the differentially expressed (DE) proteins and selected all CpGs located in the DE proteins for further analysis. Although no significant individual changes were observed, we noticed trends in some CpGs that are concordant with changes of the protein level (**Figure 5A**). To further confirm our findings, we performed additional analyses in a separate cohort of 65 participants from whom DNA methylation data was publicly available (GSE151407 & GSE171140) at baseline and after 4 weeks of exercise. We then evaluated if overall DNA methylation changes were concordant with those observed at the protein level. However this analysis is solely based on group changes as multiple timepoints were not available to estimate within-subject variability and therefore to investigate individual changes. Of 192 CpGs located on the seven selected proteins (TRIM28, USP2, RHOT, AMPD3, SOD3, EPN1 and LYRM7), 32 were significantly changed after 4 weeks of exercise (p-value <0.05, **Supplementary Table 4**, see few examples on **Figure 5B**). Suggesting that these changes are occurring at the group level and potentially would also be observed at the individual level considering a larger sample size is studied and multiple timepoints were available for analysis.

**Figure 5:**
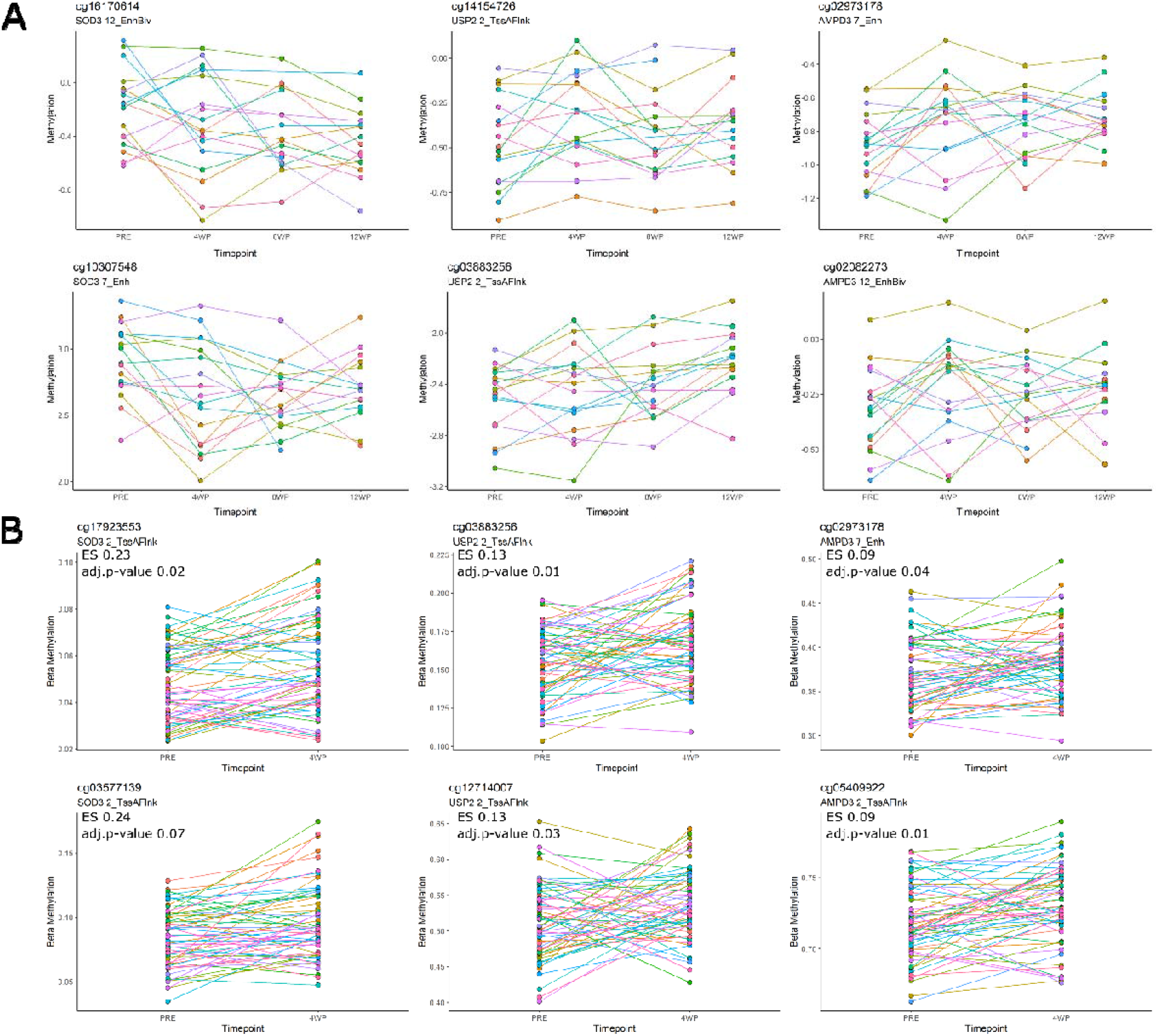
Methylation changes over time and after a 4-week HIIT intervention. A: Example of individual trajectories on few methylation sites (CpGs) located in genes from proteomic analysis presenting significant individual changes. Methylation values are based on M-values. B: Based on the same significant genes from individual analyses we have evaluated methylome changes pre/4-week post exercise in a larger cohort (GSE151407 & GSE171140). In these plots individual changes could not be estimated however we can see stronger (significant) changes to occur at the group level when sample size is increased. ES=Effect size. For both graphs A/B each participant is represented by a different colour.

Protein changes can sometimes be a reflection of changes in expression levels of mRNA in the respective gene. Thus, to investigate some of our findings at the transcriptome level we selected three of the top proteins with effect size of >0.5 (AMPD3, SOD3 and USP2) and performed real-time PCR analysis in the same participants to confirm that observed protein changes are reflected at the transcriptome level. No significant changes across time (p-value >0.05) were observed for the selected mRNAs in the group level, and individual changes were only significant (p-value=0.003) for USP2.

This is unsurprising as transcriptome changes may be occurring at different timepoints than those collected (i.e. after 1h of exercise bout, after 3h of exercise bout). Based on a comprehensive meta-analysis by Amar et al. (2021) who investigated time trajectories of exercise induced transcriptome changes in skeletal muscle [69], AMPD3 and USP2 protein expression significantly increased after an acute bout of exercise while RHOT1 expression decreased after acute exercise. After long-term exercise, the protein expression of TRIM28, RHOT1 and USP2 significantly decreases, concordant with our results. Only USP2 was significant at the individual level based on our analysis. We can only speculate that the other proteins would be true at the individual level if a larger cohort was studied as the transcriptome results derived from the extrameta tool [69] are only based on group changes.

To further investigate the relationship between the methylome and the proteome, we applied a holistic and unbiased multilevel analysis using the mixOmics package [53; 54]. The multilevel analysis accounts for variation observed between individuals with subtle differences which otherwise would be masked by individual variation. After accounting for repeated measures, we observed a clear shift from baseline up to 8WP for all individuals when methylome and proteome were integrated. This indicate the clear contribution of both omics to individual changes over time (**Figure 6A**). 261 proteins were strongly positively correlated with 45 CpGs, and the same proteins were also strongly negatively correlated with 45 different CpGs (**Figure 6B & 6C**) as shown in the middle section of **Figure 6C**. Many of the 261 proteins are related to mitochondrial functioning (i.e. UQCRFS1, IDH3B, ATP5PB, NDUFA4, etc.), mitochondrial calcium ion transport (i.e. PHB, AFG3L2, VDAC1, PHB2, VDAC2, VDAC3, LATM1), mitochondrial fatty acid beta-oxidation (i.e. ACADS, MMUT, ACAA2, ECHS1, HADHA, HADHB, etc.), mitochondrial protein import (i.e. IDH3G, TIMM44, LDHD, FH, CHCHD3, CS, etc.), respiratory electron transport (NDUFV1/2/3, NDUFA4/6/7/8/9/10/12/13, SDHA/B, COX5A/B, etc.), and metabolism pathways (i.e. GLUD1, SCO1, SDHD, UQCRQ, SLC15A1, ATP5PO, ACSL1, etc.).

**Figure 6:**
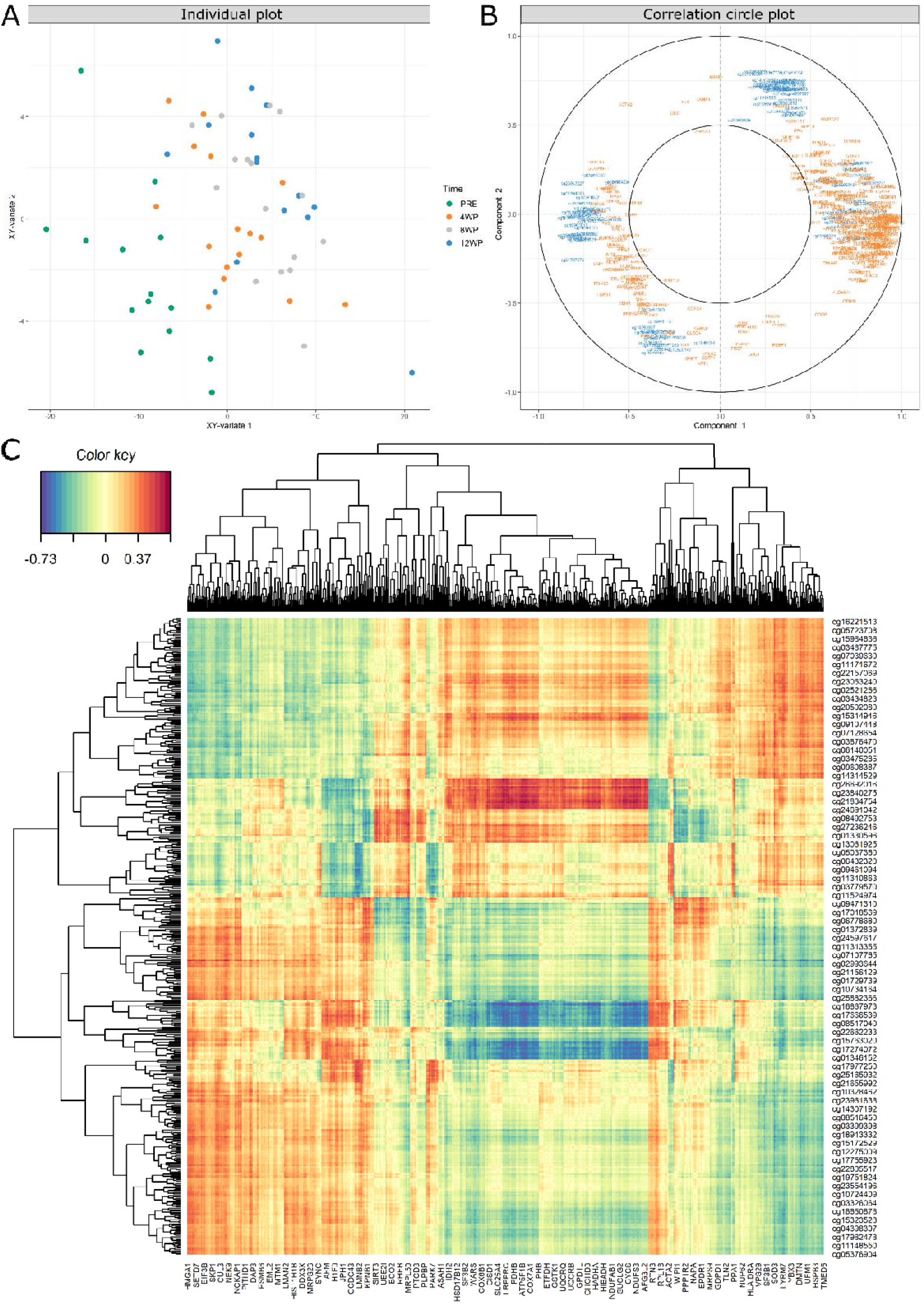
Multilevel integration of methylome and proteome. A: Individual plot based on methylome and proteome integration data after controlling for repeated measures. Samples are projected into the space spanned by the averaged components of both methylome (x) and proteome (y). Each dot represents an individual and different colors represent different timepoints. This plot shows that samples within the same timepoint tend to cluster and shift slightly over time as indicated by the colours. B: Correlation circle plot derived from the sparse Partial least Squares multivariate (sPLSm) analysis performed on the methylome (blue) and proteome (orange) data after controlling for repeated measures. This plot represents the relationship between methylome and proteome markers, were points close together represent markers that are similar to each other. Markers located on the right side of component 1 represent positive correlations between variables while those in the left represent a negative correlation between variables. This plot highlights the contributing variables that together explain the covariance between the two datasets. The middle circle represents a cutoff = 0.5 which highlights only the variables with stronger contributions to each component found in the outer circle. C: Clustered Image Map from the sPLSm after controlling for repeated measures. The plot displays the similarity values between the methylome (rows) and proteins (columns) variables selected across two dimensions, and clustered with a complete Euclidean distance method. Cell colours represent the correlation between methylome and proteome (i.e. blue negative correlation and red positive correlation).

## 4. Discussion

We found that intensified exercise training induced changes in the methylome and, to a greater extent the proteome in human skeletal muscle. Over four-hundred proteins significantly changed after exercise, and k-means analysis revealed clear clustering highlighting timely changes exhibited by each group of proteins. Utilising repeated testing approach, we uncovered over one-hundred proteins associated with individual responses to exercise, and seven proteins had a large effect size of >0.5. Only one DNA methylation loci (DMP) was significantly associated with individual responses to exercise (cg06587054 which is annotated to the *PARL* - Presenilin-associated rhomboid-like gene located on chr3:183884873-183884875).

In the group level analyses, only one DMP decreased in methylation levels after 12 weeks of exercise (cg23669611; *RTN4RL1;CTD-2545H1.2* – Reticulon 4 receptor like 1 gene located on chr17:1998389-1998391). According to the human protein atlas the protein annotated to this gene protects motor neurons against apoptosis [58]. However, its role in skeletal muscle and in response to exercise is unknown. In our proteomic analyses, RTN4RNL1 was not associated with exercise responses, however RTN4 (Reticulon 4) an associated protein (i.e. shared function) [70] significantly decreased its expression after exercise (ES: -0.07, adj.p-value: 0.05).

In the individual level analyses, we also found decreased methylation levels in one DMP annotated to the *PARL* gene (cg06587054). *PARL* has a pivotal role in the quality control and maintenance of the mitochondria steady state [55; 71]. Interestingly, *PARL* knockout mice have been shown to suffer from progressive multisystem disease characterized by a progressive muscle and immune organ atrophy [71; 72]. We report a consistent individual decrease in methylation thus, it is possible that decreased DNA methylation in this DMP following exercise provides a protective effect against atrophy. This DMP (cg06587054) is located in a CpG island in an active transcription chromatin state, thus we hypothesise that exercise may induce decrease in methylation of this gene which in turn will lead to increase gene and protein expression. Further targeted investigation in larger cohorts in all omics is warranted to confirm causality and elucidate exercise effects in skeletal muscle *PARL* expression.

We found many proteins significantly associated with exercise at both the group and individual level analyses. Group changes highlight modifications in mitochondrial related pathways with most proteins increasing in expression in an exercise dose-dependent manner, a concept that has been well described by the literature [37; 73-78]. Interestingly, we found that mitochondrial ribosomal proteins increased with exercise while nuclear ribosomal proteins decreased with exercise. Both mitochondrial and nuclear genome encode for components of the mitochondrion’s ribosome [79]. Ribosomal proteins are encoded in the nuclear genome and then imported to the mitochondria where they assemble with the 2 rRNAs to form mitochondrial ribosomes, which are responsible for translation of 13 mitochondrial mRNAs [79]. Previous study showed that ribosomes are likely to increase in response to resistance training and induce hypertrophy [80]. The exercise applied in the current study was endurance (HIIT), and is not likely to induce significant hypertrophy, hence this might be one of the explanations to the observed results. However, in depth analysis including skeletal muscle RNA content, ribosomal biogenesis and targeted comparison between mitochondrial ribosomes and nuclear ribosomes in response to different types of exercise is warrant to confirm these findings.

Overall, we identified seven proteins associated with individual responses to exercise, with two novel exercise-related proteins, *LYRM7* and *EPN1*. LYR motif containing 7 (*LYRM7*), also known as Complex III assembly factor and its deficiency has been previously associated with mitochondrial complex III defects [81]. Furthermore, low expression of the gene has also been associated with lactic acidosis, muscle weakness and exercise intolerance [82]. We found significant increased expression of LYMR7 protein in a time-dependent manner in our analyses. Thus, higher expression of *LYRM7* gene and protein may have a positive influence on skeletal muscle function, and it is candidate for functional and molecular analyses to further understand its potentially protective role in skeletal muscle. *EPN1* is a major component of clathrin-mediated endocytosis [83]. Clathrin-dependent endocytosis allows the uptake of extracellular molecules and transport it into the intracellular region within a series of vesicle compartments called endosomes [84]. Proper trafficking of such components are vital for cell growth and survival. *EPN1* is also known to interact with a transcription factor (PLZF) which in turn may influence transcription of specific genes [84]. The positive role of EPN1 protein in exercise adaptation and skeletal muscle functioning remains to be investigated.

Three (*SOD3, RHOT1* and *LYRM7*) of the seven proteins with large effect size were previously associated with exercise in tissues other than muscle, and hence may have a systemic or cross-tissues effect. Superoxide dismutase 3 (*SOD3*) is a redox enzyme that reduces the potential toxicity of reactive oxygen species induced cellular damage [85]. Increased expression of *SOD3* after exercise was reported in plasma exosomes, in both mice and healthy humans [61]. In mice, *SOD3* overexpression in skeletal muscle is released to circulation to exert cardio protective effects [61]. Here, we observed a steady increase in *SOD3* protein expression after exercise in all the study participants, suggesting that similar mechanisms may be observed in humans as those reported in mice. To prove the theory, future, targeted studies should measure plasma exosomes, and investigate if and how *SOD3* mediate exercise-induced angiogenesis. The Ras homolog family member T1 (*RHOT1*) is a mitochondrial GTPase involved in mitochondrial trafficking [86], and has been reported to decreased its expression after exercise, in rat liver [62]. In the present study, we showed decreased *RHOT1* expression in skeletal muscle following exercise. The mechanism by which RHOT1 is influencing skeletal muscle adaptation to exercise should be explored further in future studies.

A recent paper investigating the effects of training in the human proteome and acetylome identified novel exercise-training regulated proteins including glutaminyl-tRNA synthase (QARS) and rab GDP dissociation inhibitor alpha (GDI1) [87]. These proteins are associated with insulin-stimulated glucose uptake and may provide a novel mechanism explaining the role of exercise training in insulin-sensitizing effects. Interestingly in our analysis both proteins presented significant changes in the same direction as their paper described (QARS - ES:0.04, adj.p-value: 0.05, GDI1 - ES:-0.06, adj.p-value:0.013), validating their findings and highlighting the potential importance that these proteins may have in insulin regulation following exercise. Future functional validation investigating such mechanisms are warranted.

We applied unbiased integration approach to uncover subtle difference in exercise responses across multiple layers (epigenome and proteome). The integration analysis highlighted proteins both positively and negatively associated with specific CpGs, worthy following up in future replication/mechanistic studies. Interestingly of the 261 proteins highlighted in the results, we found proteins positively correlated with CpGs located in genes involved in fatty acid metabolism such as *COMMD9*, (involved in LDL regulation) [88]; hypertension (*ARHGAP42)* [89]; weight loss and muscle development (*APOBEC1*) [90]; obesity (*PTPRT* and *PTPRN2)* [91-94], and muscle myogenesis and hyperthrophy (*DACT1, DIO2, RPTOR*, and *PLEKHM3)* [95-99] These proteins were also negatively correlated with CpGs located in genes such as *KCNMA1, NOTCH1*, and *CAMKK2* which are involved in skeletal muscle regeneration, proliferation and differentiation [100-105], *DHRS3, DGKG, LPIN1, WNT5A* which have been associated with obesity, metabolic syndrome and lipid regulation [106-108], *LRRC2* which is a mediator of mitochondrial and cardiac function [109], and *PDE4A* which has been associated with diabetes [110; 111]. Based on results of this integration, we suggest that future work studying the relationship between such genes/proteins should be prioritized as this may reveal important mechanisms associated with exercise response across multiple OMIC layers.

## 5. Conclusions

Collectively, we found a significant influence of exercise in multiple omic layers, with a strong effect of high-intensity exercise on the proteome, in a dose-dependent manner. Novel proteins, that have not been previously associated with either skeletal muscle or exercise response, consistently changed in the group level and across individuals (individual response), highlighting the robust effect these proteins have on exercise responses. We note that a limitation of this study is the relatively small sample size masking some of the effects the methylome has on exercise. Furthermore, the effects of exercise is systematic. In fact some proteins observed in our study may affect other tissues. Hence, systemic investigations including multiple tissue-types following exercise will help to shed light into the benefits that exercise exerts in the human. Future mechanistic and validation studies are required to validate our novel findings.

## Acknowledgements

We would like to express gratitude to all of our participants and their efforts during the intervention making this study possible. This work was supported by Nir Eynon’s NHMRC Career Development Fellowship (APP1140644), and NHMRC Investigator Grant (APP1194159). The Gene SMART study is also supported by an Australian Research Council (ARC) Discovery Project Grant (DP190103081). This study used BPA-enabled (Bioplatforms Australia) / NCRIS-enabled (National Collaborative Research Infrastructure Strategy) infrastructure located at the Monash Proteomics and Metabolomics Facility.

